# Single-cell RNA sequencing reveals characteristics of myeloid cells in pulmonary post-acute sequelae of SARS-CoV-2

**DOI:** 10.1101/2023.07.31.551349

**Authors:** Hyundong Yoon, Logan S. Dean, Boonyanudh Jiyarom, Vedbar Khadka, Youping Deng, Vivek R. Nerurukar, Dominic C. Chow, Cecilia M. Shikuma, Gehan Devendra, Youngil Koh, Juwon Park

## Abstract

**Background:** Although our understanding of the immunopathology and subsequent risk and severity of COVID-19 disease is evolving, a detailed account of immune responses that contribute to the long-term consequences of pulmonary complication in COVID-19 infection remain unclear. Few studies have detailed the immune and cytokine profiles associated with post-acute sequalae of SARS-CoV-2 infection with persistent pulmonary symptoms (PPASC). However, the dysregulation of the immune system that drives pulmonary sequelae in COVID-19 survivors and PASC sufferers remains largely unknown.

**Results:** To characterize the immunological features of pulmonary PASC (PPASC), we performed droplet-based single-cell RNA sequencing to study the transcriptomic profiles of peripheral blood mononuclear cells (PBMCs) from participants naïve to SARS-CoV-2 (Control) and infected with SARS-CoV-2 with chronic pulmonary symptoms (PPASC). We analyzed more than 34,139 PBMCs by integrating our dataset with previously reported control datasets (GSM4509024) cell distribution. In total, 11 distinct cell populations were identified based on the expression of canonical markers. The proportion of myeloid-lineage cells ([MLCs]; CD14^+^/CD16^+^monocytes and dendritic cells) was increased in PPASC compared to controls. MLCs from PPASC displayed up-regulation of genes associated with pulmonary symptoms/fibrosis, while glycolysis metabolism-related genes were downregulated. Similarly, pathway analysis showed that fibrosis-related (*VEGF*, *WNT*, and *SMAD*) and cell death pathways were up-regulated, but immune pathways were down-regulated in PPASC. In PPASC, we observed interactive *VEGF* ligand-receptor pairs among MLCs, and network modules in CD14^+^ (cluster 4) and CD16^+^ (Cluster 5) monocytes displayed a significant enrichment for biological pathways linked to adverse COVID- 19 outcomes, fibrosis, and angiogenesis. Further analysis revealed a distinct metabolic alteration in MLCs with a down-regulation of glycolysis/gluconeogenesis in PPASC compared to SARS-CoV-2 naïve samples.

**Conclusion:** This study offers valuable insights into the immune response and cellular landscape in PPASC. The presence of elevated MLC levels and their corresponding gene signatures associated with fibrosis, immune response suppression, and altered metabolic states suggests their potential role as a driver of PPASC.

## Background

Three years have passed since the outbreak of the novel coronavirus disease 19 [COVID-19], caused by severe acute respiratory syndrome virus 2 [SARS-CoV-2] (1). COVID-19 is a global public health crisis and has had profound impacts on economic growth and social structures (2–4). The advent of COVID-19 vaccines has effectively reduced the risk of SARS-CoV-2 infection and prevented death and severe illness following acute infection (5, 6). However, approximately 30- 50% of COVID-19 survivors experience chronic health conditions (‘sequelae’) that persist longer than 3 months and develop within 28 days of acute illness (7–9). These ongoing health problems are termed Long COVID but also referred to as “post-acute sequelae of SARS-CoV-2” (PASC). PASC symptoms reported from survivors varies, but common symptoms include fatigue, insomnia, dyspnea, cough (10). A subset of PASC is termed pulmonary PASC (PPASC), due to the chronic respiratory nature of the long-term symptoms (10). Recent large cohort studies demonstrated that COVID-19 survivors have a greater risk of developing comorbidities, particularly diabetes (11), as well as cardiovascular (12) and kidney (13) diseases. Considering the global prevalence of PASC cases (14–16) and the impact of sequelae on COVID-19 survivors’ quality of life (17–19), PASC continues to present a significant global burden to healthcare infrastructure.

The etiology of PASC, and therefore PPASC, is not yet known, but ongoing studies have provided risk factors and potential predictors of PASC development. Factors, such as female sex, age, comorbidities, smoking, and social determinants of health (low socioeconomic status, and racial/ethnic minority) are associated with an increased risk of PASC development (20–22). While severity of COVID-19 severity is suspected to be a risk factor for PASC (23), evidence continues to present mixed conclusions (24).

Investigations into the immunopathology associated with PASC have revealed that aberrant cellular and humoral immune responses are drivers of PASC (25–28). Comparison of the circulating proteome between individuals with acute COVID-19, COVID-19-recovered and PASC demonstrate no significant differences in most cytokines related to inflammation, nor a disease-specific signature among the groups (25, 29). However, a consensus of studies has demonstrated IL-6, TNF-α, and IL-1β are commonly elevated in individuals with PASC (30–32), suggesting a pro-inflammatory etiology.

Cellular metabolic dysregulation is one of the major features of SARS-CoV-2 infection and a key determinant of disease severity (33, 34). Furthermore, epidemiological observations demonstrated that individuals who had metabolic comorbidities, such as diabetes mellitus, hyperglycemia, or obesity were at a significantly increased risk of severe COVID-19 (35). Lung epithelial cells and monocytes infected with SARS-CoV-2 demonstrate lipid dysregulation with excessive lipid droplet accumulation within the cells (36–38). Inhibition of lipid droplet biogenesis was demonstrated to inhibit the proinflammatory and replication of SARS-CoV-2 within infected monocytes (36), suggesting a metabolic rearrangement towards lipid use for viral replication. Investigations into glycolysis metabolic pathways reveals a strong propensity for upregulation within macrophages, monocytes, and T cells, consistent with altered cell activation and attenuated T cells’ effector functionality (39, 40). Recent evidence demonstrates a metabolic shift towards a pro-resolution and remodeling phenotype in macrophages isolated from PASC patients (41), indicating changes in plasma metabolites after SARS-CoV-2 infection led to profound and prolonged cellular metabolic disruption and subsequent cell dysfunction.

The persistence of SARS-CoV-2 protein and RNA in both immune cells and tissues (42, 43) have arisen as potential factors for PASC development (44). Persistence of viral components is well documented to have prolonged impacts on immune cell functionality and polarization (45–47), and therefore likely influence PASC etiology. It is likely a combination of a multitude of the aforementioned factors driving PASC development. As such, each avenue of investigation remains a viable option for discovery, with the goal of predictive and/or therapeutic intervention innovations for PASC.

Recent single-cell RNA sequencing (scRNA-seq) data has demonstrated profound phenotypic alteration of immune cells in COVID-19 convalescents with persistent symptoms. Innate and adaptive immune cell perturbations are evident by 12 weeks post infection and sustained within individuals who develop PASC (48). Another large-scale single cell sequencing endeavor revealed the predictive value of cortisol levels as well as circadian rhythm-related gene signatures as predictive of PASC development (49). Although ongoing PASC studies are focused on advancing our understanding of its’ pathophysiology, the mechanisms underlying persistent pulmonary sequelae secondary to SARS-CoV-2 infection remain largely unexplored. The manner in which key immune cell subsets change their functionalities in pulmonary PPASC have remained undefined. Therefore, identification of primary cell types associated with PPASC and signaling pathways mediated by immune cell alteration and activation are crucial to obtaining insights into immune cell perturbation that contribute to prolonged pulmonary sequelae in PASC.

In this study, we analyzed PBMCs from individuals who had developed PPASC after SARS-CoV- 2 infection and uninfected controls. The immune profiling and transcriptomic data revealed alterations of myeloid lineage immune cells, notably monocytes, and dendritic cell in PPASC. We further explored individual cell-cell interactions on immune cell types and performed metabolic analyses, demonstrating a profound association with fibrosis, vascularization, and characteristic of a reparative immune environment. Together, our results provide evidence that perturbations undergone within the myeloid lineage of immune cells are likely associated with ongoing pro-fibrotic processes that may therefore be contributing specifically to PPASC development. This provides valuable insight into the utility of targeting immune cell populations to ameliorate pulmonary sequelae among PPASC individuals.

## Materials and Methods

### Peripheral blood mononuclear cell isolation

Human venous peripheral blood samples were taken at the state’s main tertiary care hospital- Ambulatory Post- COVID Clinic (Queen’s Medical Center, Honolulu, Hawaii) and collected in ethylene diamine tetra-acetic acid (EDTA) tubes (BD, Vacutainer) by venipuncture. In brief, venous blood was diluted with an equal volume of phosphate buffered saline (PBS) and layered on top of Ficoll-Paque Plus (GE Healthcare Biosciences, Piscataway, NJ) following the manufacturer’s protocol. Peripheral blood mononuclear cells (PBMC) were separated by centrifugation at 400 × g for 30 minutes at room temperature (RT). PBMC were collected from the buffy coat, red blood cells lysed, and the remaining pellet washed twice in PBS supplemented with 2% fetal bovine serum (FBS). Cells were then counted, viability determined, and cryopreserved at 5 million cells/vial.

### Pulmonary function tests

Pulmonary function testing (PFT) was performed on individuals with Pulmonary PASC (PPASC). All PG participants underwent PFTs (Vyaire) with the measurements of forced vital capacity (FVC), forced expiratory volume in 1 second (FEV1), total lung capacity (TLC), and DLCOc% interpreted in accordance with European Respiratory Society (ERS)/American Thoracic Society (ATS) guidelines (50).

### Fluorescence Assisted Cell Sorting

For live PBMC purification for single cell RNA sequencing, PBMC were stained with BV711- CD45 (1:200 dilution, Biolegend, San Diego, CA), for 30 minutes at RT after adding Human TruStain FcX (1:200 dilution, Biolegend, San Diego, CA) for 5 minutes. Propidium iodide (PI) was added just before cell acquisition to assess cell viability. UltraComp eBeads Compensation Beads (Thermo Fisher Scientific, 01-2222-42) were used for compensation. Approximately 98- 99% of total PBMCs were identified as live, live CD45^+^ cells were sorted into 0.1% bovine serum albumin (BSA) in DPBS by a FACSAria IIu Cell Sorter (BD Biosciences, Franklin Lakes, NJ). Sorted cell suspension were pelleted at 350 x g for 5 minutes at 4 °C, and resuspended in 1% BSA in DPBS to about 1,000 cells/μl.

### Single-cell RNA-Sequencing

Cell concentration and viability were confirmed using automated cell counter Countess II (ThermoFisher Scientific) with 0.4% trypan blue solution and samples with viability > 70% were further processed. To obtain single-cell gel beads-in-emulsion (GEM), cell suspension was pelleted by 400 x g for 5 minutes at 4 °C, resuspended cells at a concentration of 1,000 cells/μl in 0.1% BSA in DPBS. GEMs were generated in Chromium Controller by combining barcoded Single Cell VDJ 5’ Gel Beads v1.1, a Master Mix with mixture of single cells, and Partitioning Oil on Chromium Next GEM Chip G. To achieve single cell resolution, cells were delivered at a limiting dilution, such that the majority (∼90 – 99%) of generated GEMs contains no cell, while the remainder largely contain a single cell.

Immediately following GEM generation, the Gel Bead was dissolved and any co-partitioned cell was lysed. Oligonucleotides containing (i) an Illumina R1 sequence (read 1 sequencing primer), (ii) a 16 nt 10x Barcode, (iii) a 10 nt unique molecular identifier (UMI), and (iv) 13 nt template switch oligo (TSO) were released and mixed with the cell lysate and a Master Mix containing reverse transcription (RT) reagents and poly(dT) RT primers. Incubation of the GEMs produced 10x Barcoded, full-length cDNA from poly-adenylated mRNA GEMs were broken and pooled after GEM-RT reaction mixtures are recovered. Silane magnetic beads were used to purify the 10x Barcoded first-strand cDNA from the post GEM-RT reaction mixture, which includes leftover biochemical reagents and primers. After cleanup 10x Barcoded, full-length cDNA was amplified via PCR with primers against common 5’ and 3’ ends added during GEM-RT. Amplification generated sufficient material to construct 5’ Gene Expression libraries. Enzymatic fragmentation and size selection were used to optimize the cDNA amplicon size prior to 5’ Gene Expression library construction. P5, P7, a sample index, and Illumina R2 sequence (read 2 primer sequence) were added via End Repair, A-tailing, Adaptor Ligation, and Sample Index PCR. The final libraries contain the P5 and P7 priming sites used in Illumina sequencers. Library quality and size distribution was confirmed on Agilent Bioanalyzer High Sensitivity DNA chips (Agilent). Library quantification was performed with qPCR using KAPA Library Quantification Kit for Illumina Platforms (KAPA/Roche). Libraries were normalized, denatured and diluted to obtain 1.5 pM molarity. Libraries were sequenced on NextSeq500 instrument at depth of minimum 20,000 reads/cell using 26 x 91 bp read settings.

### Processing and analysis of scRNA-seq data

The scRNA-seq raw fastq files were aligned to the human reference genome (ver. GRch38) using the Cell Ranger pipeline (51) to generate a raw gene-by-cell count matrix. Empty droplets were identified (52) and real cell data satisfying FDR < 0.01 were retained. Low-quality cells were removed (53) based on overall cell quality calculation. Outliers were identified using principal component analysis (PCA) and removed, and cell-by-cell bias was removed by clustering cells (54). Cell-specific size factors were calculated. Gene-by-cell matrix normalization was performed by dividing the number of raw unique molecular identifiers (UMIs) by the cell-specific size factor. Normalized counts were log2 transformed by adding a pseudo count of 1. One thousand highly variable genes (HVGs) for biological variability were selected. Using the k-nearest neighbor (kNN) graph was calculated based on different principal components (PCs) in each sample, and cell clusters were visualized using uniform manifold approximation and projection (UMAP).

### Single cell RNA data merge and integration analysis

To integrate scRNA data generated from different batches, Harmony (55) was used to mitigate batch effects by merging and integrating the data.

### Cell type identification

Investigate the differential expression of well-known canonical marker genes in each cluster. Additionally, cell types were identified by visually checking the average expression values of canonical marker genes (56).

### Cell proportion test

Confirmed the cell proportion by counting the number of cells of each cell type as a percentage. A permutation test between groups was conducted (57) to calculate the p-value/ false discovery rate (FDR). Finally, we performed bootstrapping to validate the difference in proportion, and the results were classified as the statistically significant intervals of Log2FD > 0.3, < 0.3, and FDR < 0.05.

### Differentially expressed gene (DEG) analysis

To assess the differential gene expression in the scRNA data of two groups (each PPASC and Control), we conducted differential expressed gene analysis with the built-in framework of MAST (58). In the analysis results, significant up- and down-regulated genes were classified based on statistical criteria of avg_log2FC > 0.25, < -0.25 and p-value < 0.05.

### Gene set enrichment analysis (GSEA)

To conduct the analysis, Hallmark gene sets (H), Curated gene sets (C2), and Ontology gene sets (C5) databases of MsigDB (59) selectively used and analyzed (60). Pathways that were significantly up- or down enriched were classified based on the Normalized Enrichment Score (NES) > 0, < 0 and p-value < 0.05.

### Cell to cell interactions

The human ligand-receptor (only Secreted signaling) database were used to analyze the ligand-receptor interaction between cells (61). Cells expressing less than 5% of genes were excluded, and only interactions with statistical significance (p-value < 0.05) among autocrine or paracrine interactions were classified. And to compare the interaction signals between the two groups, the communication probability value calculated based on the geometrical mean expression level was confirmed.

### Pseudo-bulk differentially expressed gene (DEG) analysis

To make single-cell RNA into pseudo-bulk RNA, we aggregated transcript count tables of all single-cells per sample to verify possible false positives in single-cell DEGs. Differential expressed analysis was performed (62), and genes satisfying logFC > 0.25 or < -0.25, along with a p-value < 0.05 were classified as up- or down-regulated genes, respectively.

### Single-cell correlation network analysis

Integrated scRNA data was utilized to calculate Pearson correlation values (63). Subsequently, correlation networks were created for specific cell types, and the identification and validation of differentially expressed network modules were accomplished through enrichment analysis (64).

## Results

### Alteration of myeloid-lineage cells in individuals with PPASC

Circulating immune cell levels and their functional status hold promise as biomarkers for assessing the severity of COVID-19. Therefore, analysis of blood immune cells in COVID-19 convalescents provides insight into the long-term consequences of host immune responses after SARS-CoV-2 infection. To explore immune system alteration associated with persistent lung sequelae after recovery of acute illness of SARS-CoV-2, we selected two participants who had reduced diffusion capacity for carbon monoxide (corrected for hemoglobin-DLCOc, <80%) by pulmonary function test (PFT) among individuals with PPASC. They had persistent pulmonary symptoms greater than 5 months after onset of SARS-CoV-2 infection. (Supplementary Table 1). As a comparator group, a participant naïve to SARS-CoV-2 infection was selected and confirmed negative via a SARS-CoV-2 nucleocapsid antibody test (Supplementary Table 1). Live CD45^+^ cells were sorted by fluorescence-assisted cell sorting (FACS) and subjected to scRNA-seq utilizing the 10X Genomic Chromium system. We sequenced 30,000 cells from three PBMC samples. To match number of control samples, we confirmed the utility of scRNA-seq data of SARS-CoV-2 naïve PBMC from publicly available scRNA-seq data of PBMC (GSM4509024)(65) to integrate our dataset. scRNA seq data of PBMC from three participants were combined into an integrated dataset of one single cell dataset of PBMC from a naïve participant (age-matched) and the similarity of gene expression profiles measured in each cell was calculated mitigating technical differences between different batches in each. Total 34,139 cells passed quality control (QC; Removing empty droplets, cells with low UMI / feature counts, high mitochondrial gene expression indicative of dying cells; Low-quality cells were filtered out by removing cells 5,861 cells, Supplementary Data Fig. 1).

To compare scRNA-seq data between groups, we subjected four independent scRNA-seq data sets with 34,139 cells to uniform manifold approximation and projection (UMAP) based on the RNA expression. Canonical marker genes of known cell types were enriched in each cluster (Fig. 1C) and the marker expressions were used to annotate the clusters. A total of 11 cell types were identified, including CD4^+^/CD8^+^ T cells, natural killer (NK) and natural killer T (NKT) cells, CD14^+^ and CD16^+^ monocytes, dendritic cells, platelets, hematopoietic stem cells (HSC), B cells, and erythrocytes (Fig. 1A). All identified immune cell populations were present in both SARS-CoV-2 infected and naïve participants (Fig. 1B). We further attempted to determine a vital feature for reflecting immune alterations associated with PPASC and the relative proportions of peripheral immune cells were examined between PPASC and control. Among the immune cells, we observed significant increases in myeloid cells (CD14^+^/CD16^+^ monocytes, dendritic cells, and NKT cells) and B cells whereas the proportions of NK cells, HSC, and erythrocytes were significantly decreased in PPASC compared to control (Fig. 1D-E).

**Figure 1:**
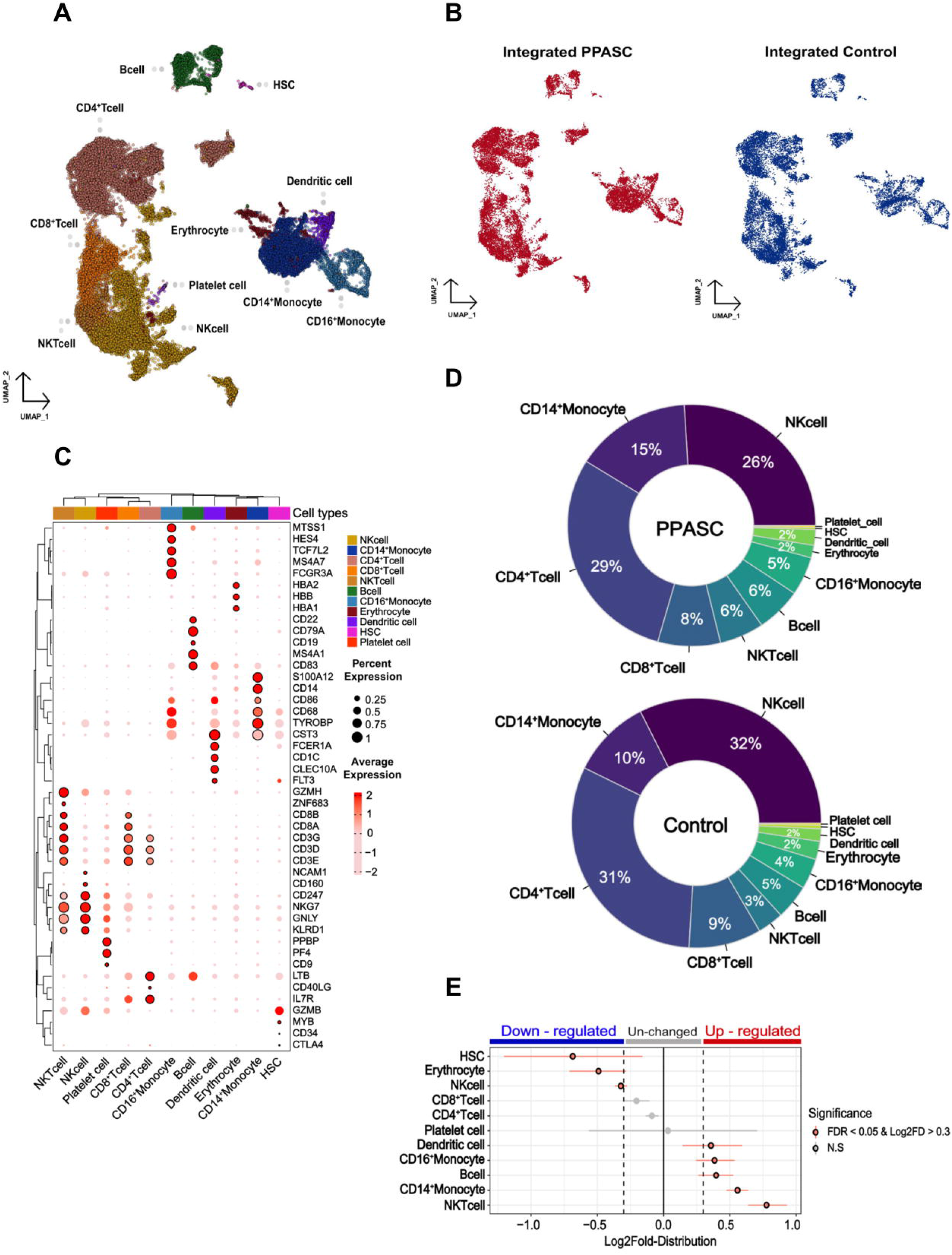
**A.** scRNA data obtained from PBMCs with a total of 34,319 integrated cells within PPASC and Controls, annotated by cell type. **B.** Distributions of single cell clusters of PPASC and Controls, respectively. **C.** Expression of canonical marker genes used for cell type identification in each cluster. **D.** Proportion of each cell type in PPASC and Control. **E.** Statistically significant up and down-regulated cell type proportions in PPASC and Control groups.

### Identification of dysregulated genes and pathways related to PPASC

To explore the characteristics of immune cells and identify the molecular changes associated with PPASC, we performed a detailed analysis of the differentially expressed genes (DEGs) of immune cells from PPASC compared with those from the control. Our scRNA-seq analysis identified pronounced alterations in myeloid cells. Furthermore, we found that elevated circulating monocyte levels and their activation were observed in COVID-19 convalescents (66). Thus, we focused our attention on myeloid cells, particularly CD14^+^/CD16^+^ monocytes and dendritic cells for downstream analysis hereafter referred to as myeloid-lineage cells (MLCs). We found a total of 872 DEGs in monocytes and dendritic cells; CD14^+^ monocytes (228 upregulated and 132 downregulated), CD16^+^ monocytes (303 upregulated and 226 downregulated), and dendritic cells (332 upregulated and 217 downregulated) (Fig. 2A-B). In addition, we demonstrate shared DEGs across three cell types (118 upregulated and 54 down-regulated,) (Fig 2A-B). Interestingly, our data showed that genes known to be associated with pulmonary symptoms/fibrosis (*ANKRD11, CTNNB1, CXCR4, HIF1A, HMGB1, ITSN2, LITAF, NEAT1, VEGFA,* and *DSE*) were upregulated in CD14^+^/CD16^+^ monocytes and dendritic cells. However, genes associated with glycolysis metabolism (*ACTB, PGK1, TPI1,* and *CFP*) were downregulated in these cell population (Fig 2A- B).

**Figure 2:**
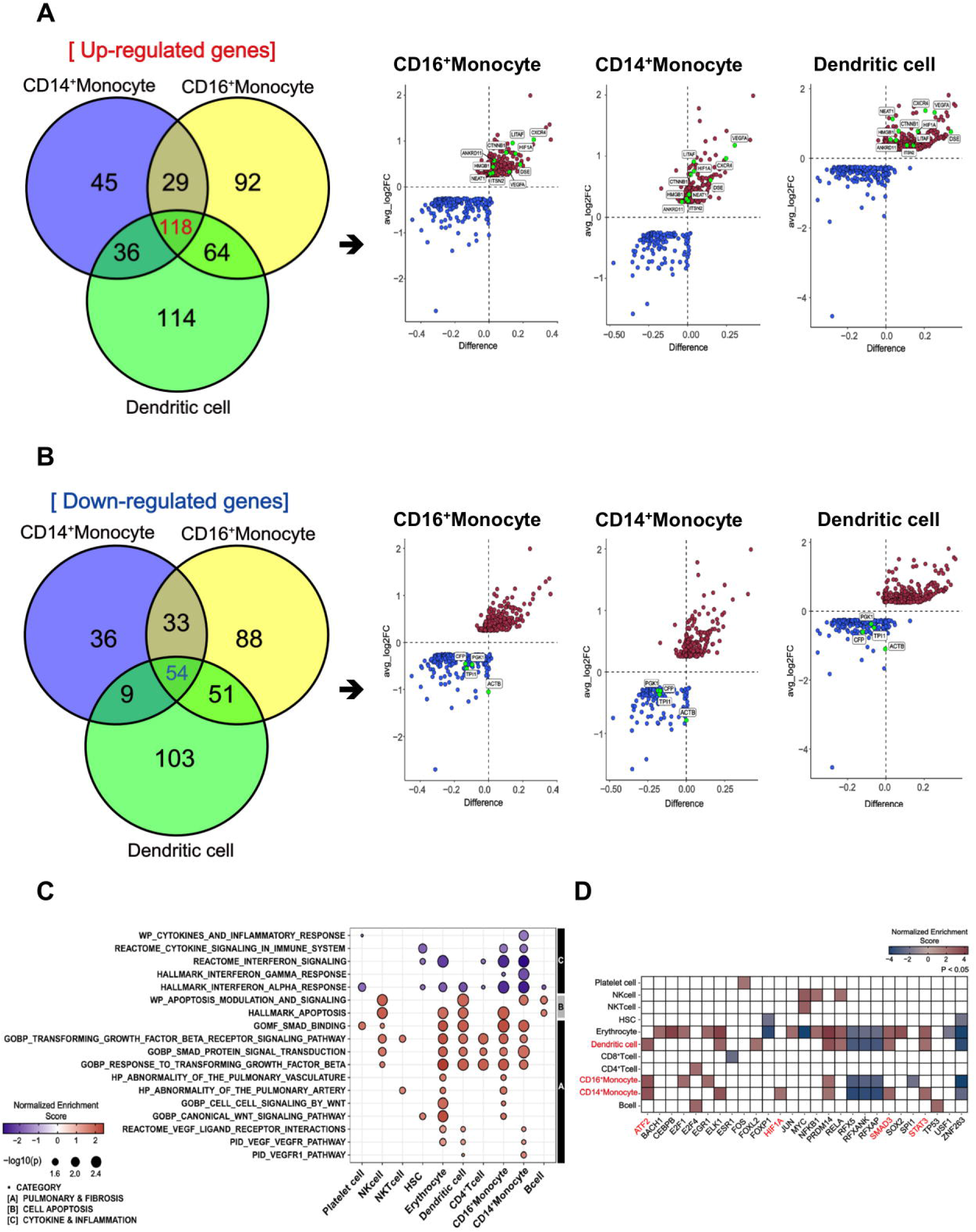
**A.** Differentially expressed and upregulated fibrosis-related genes shared between MLCs. **B.** Differentially expressed and downregulated glycolysis-related genes shared between MLCs. **C.** Enriched pathways between PPASC and Control groups in whole cell types. **D.** Statistically significant inferences of differentially activated transcription factors between PPASC and Control groups in whole cell types based on target genes expression.

Next, we performed gene set enrichment analysis (GSEA) across each cell type and conducted direct comparisons of the PPASC versus the control groups, respectively. Overall, compared to control, PPASC displayed upregulated pathways related to pulmonary and fibrosis, such as *VEGF*, *WNT,* and *TGF-β* and cell apoptosis while pathways related to cytokine/inflammation (including interferon) were downregulated in CD14^+^/CD16^+^ monocytes and dendritic cells (Fig 2C). The *VEGF* pathway was upregulated in CD14^+^ monocytes and dendritic cells, while the *WNT* pathway was upregulated only in CD16^+^ monocytes and NKT cells among all immune cell population (Fig 2C).

To identify potential direct target involved in the induction of profibrotic features in myeloid cells, we performed transcription factor enrichment analysis (TFEA) to identify transcription factors inferred from the perturbation of gene signatures between PPASC and control. We found that a total of 28 transcription factors were significantly dysregulated in individual cell types between groups (Fig 2D). Consistent with earlier analyses, we found that the downstream transcription factors of *VEGF* (*HIF1A*; CD14^+^ monocytes), *TGF-β* (*SMAD* and *STAT3*; CD14^+^ monocytes and dendritic cells, ATF2; CD14^+^/CD16^+^ monocytes and dendritic cells) signaling was upregulated in MLCs (Fig 2D). These observations demonstrate that PPASC display enriched genes driving pulmonary symptoms and profibrotic features in MLCs and that increased MLCs proportions and the altered gene signatures likely contribute to development of pulmonary sequelae after SARS-CoV-2 infection.

### Gene modules predict variable expressions and functionalities of monocytes in PPASC

To gain a comprehensive view on how the inferred targets may interact regarding PPASC development, we then identified potential cell–cell interactions and differential ligand-receptor interactions that are conserved in PPASC. The pathway of *VEGF* interaction (*VEGFA-FLT1/KDR*) was increased among CD14^+^/CD16^+^ monocytes and dendritic cells, compared to controls (Fig. 3A). We observed high fidelity communications via *VEGF* pathways in CD14^+^/CD16^+^ monocytes and dendritic cells via an autocrine or paracrine manner (Fig. 3A). Furthermore, using pseudo- bulk single cell data, we validated expression of genes identified from ligand-receptor interaction analysis. Notably, *VEGFΑ* was highly expressed in CD14^+^ and CD16^+^ monocytes, as well as dendritic cells (Fig. 3B, C).

**Figure 3:**
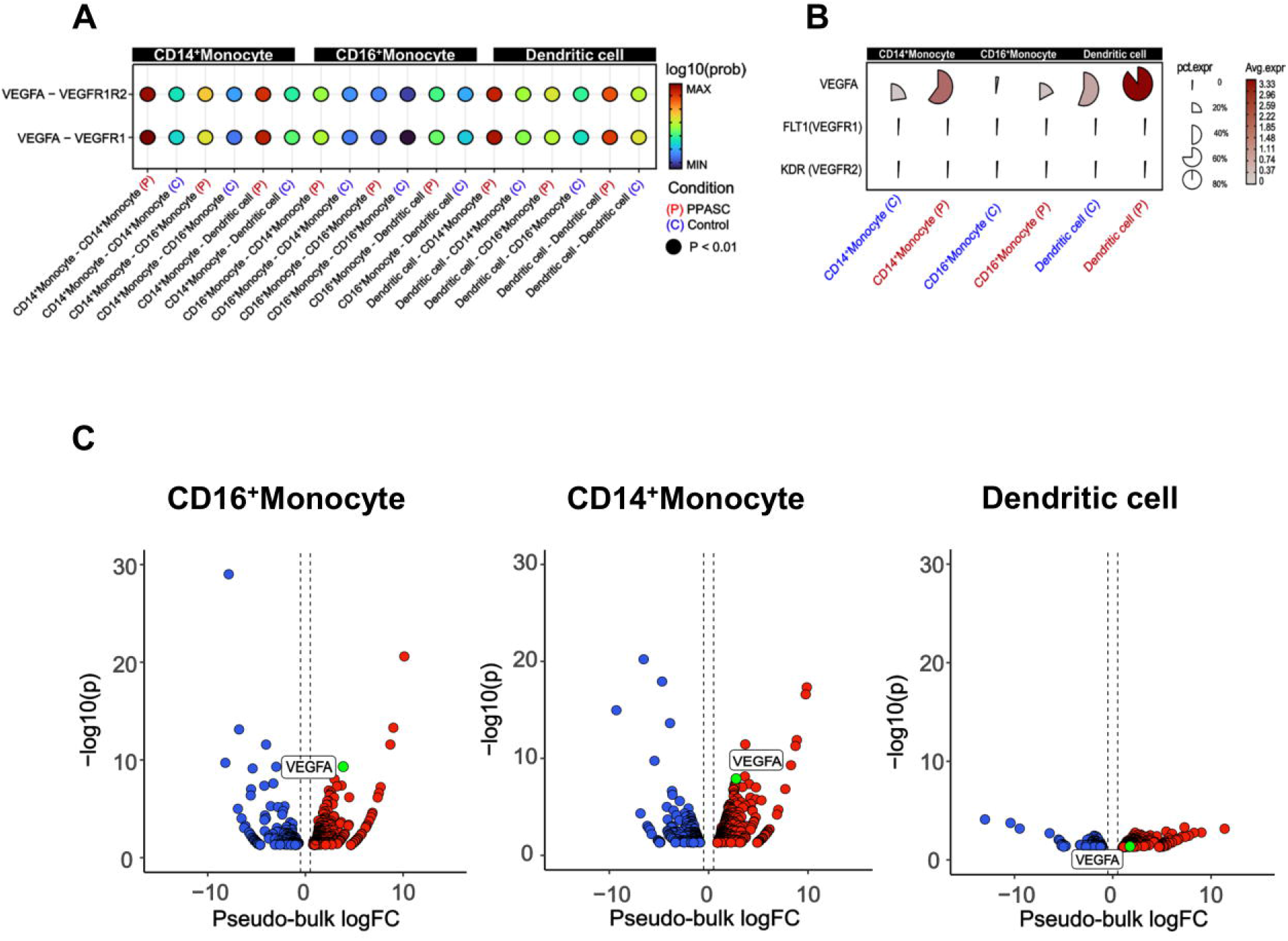
**A.** VEGF and TGF-β signaling pathway interactions between myeloid cells in PPASC and control groups. **B.** Average and percent expression of ligand-receptor genes in myeloid cells in PPASC and control groups. **C.** Differentially expressed ligand genes in aggregated and pseudo- bulked single cells.

Within the monocyte clusters (CD14^+^ and CD16^+^ monocytes), we further sub-grouped the signatures into gene-correlation based on network modules (Fig 4A). Within CD14^+^ monocyte populations, network module 5 (Net-M5) was specifically found to be upregulated, while CD16^+^ monocyte demonstrated an upregulation of network module 4 (Net-M4) (Fig. 4B). To identify the organizing hub genes in Net-M4 and M5, the top 60 genes with high eigengene-based connectivity (kME) values were selected (Fig. 4C). In the Net-M4, genes involved in *VEGFA-VEGFR2* signaling (*NR4A1, TPM3, FAM120A*, and *NUMB*) and *WNT* signaling (*HHEX* and *APC*) were identified. The Net-M5 detected genes involved in lung fibrosis (*CXCL8* and *IL1B*), *VEGFA- VEGFR2* signaling (*NR4A1* and *CBL*), and *TGF-β* signaling (*JUNB* and *ATF3*) (Fig, 4C). Furthermore, the pathway enrichment analysis of Net-M4 and Net-M5 revealed that CD16^+^ M4 module was associated with *WNT*, *VEGF*, and *TGF-β* signaling pathways (Fig. 4D). The CD14^+^ M5 module was associated with COVID-19 adverse outcome linked pathways as well as fibroblast growth factor stimulation pathways. Interestingly, this same module in CD14^+^ monocytes revealed increases in pathways for fibroblast-specific apoptosis and general apoptotic pathways (Fig. 4D).

**Figure 4:**
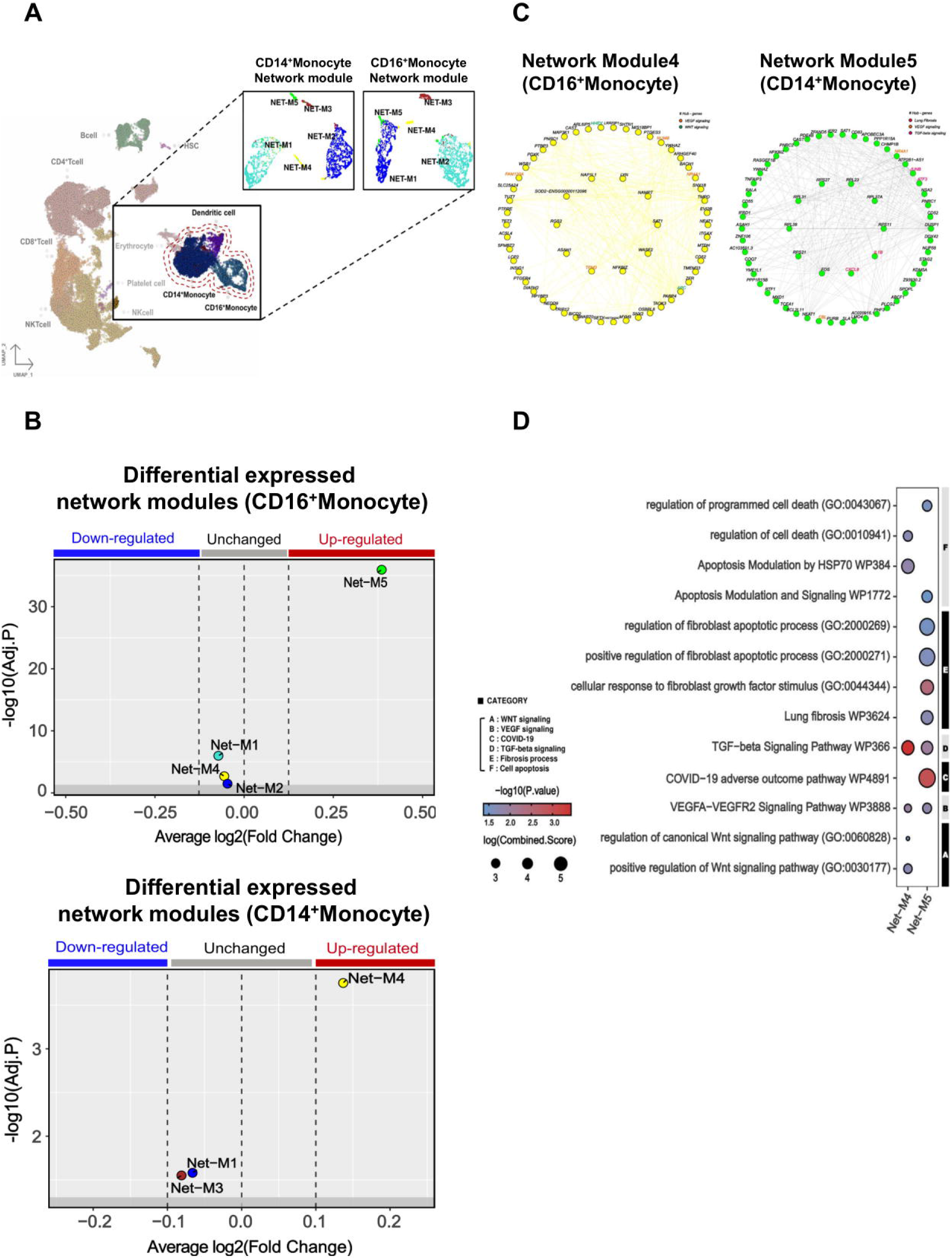
**A.** Correlation-based network module in CD14^+^ and CD16^+^ monocytes incorporating PPASC and control groups. **B.** Differentially expressed network modules in CD14^+^ and CD16+ monocytes in PPASC and controls. **C.** Hub genes of the M4 and M5 from the differentially expressed network modules. **D.** Enriched pathways in the differentially expressed network module.

### PPASC display metabolic alterations in myeloid subsets

Reports of the metabolic impacts of SARS-CoV-2 infection and association with chronic fatigue and diabetic-like events during Long-COVID are well documented (67). These observations aligned with our findings of downregulated glycolytic metabolism and prompted us to determine the impact of SARS-CoV-2 infection on immunometabolism and its relevance to pulmonary sequelae (Fig. 5A). Using Gene Set Variation Analysis (GSVA) based on non-parametric unsupervised learning to infer the variability of gene set enrichment related to gluconeogenesis/glycolysis, we observed a decreased average NES score in MLCs of PPASC compared to the control group. The inferred metabolic state of individual cells using a database of integrated metabolic networks and metabolic flux balance analysis demonstrated that MLCs of PPASC exhibited downregulation of gluconeogenesis/glycolysis metabolic reactions compared to the control group (Fig. 5B). Furthermore, inference of intercellular metabolite communication based on estimated metabolite scores and averaged sensor gene expression values by cell group demonstrated that the metabolite, D-Glucose, was directly associated with gluconeogenesis/glycolysis metabolism. Furthermore, these inferred interaction scores revealed decreased mean abundance values in MLCs of PASC compared to the control group (Figure 5C). The calculated communication scores for metabolite-sensor partners revealed statistically significant interaction scores in MLCs of the control group but not in PPASC (Fig. 5D). This suggests a lack of gluconeogenesis/glycolysis-related metabolite interaction or a diminished interaction in PPASC. In summary, these metabolic analysis observations indicate a decreased regulation of the previously identified glucose-specific metabolic pathway in MLCs of PPASC.

**Figure 5:**
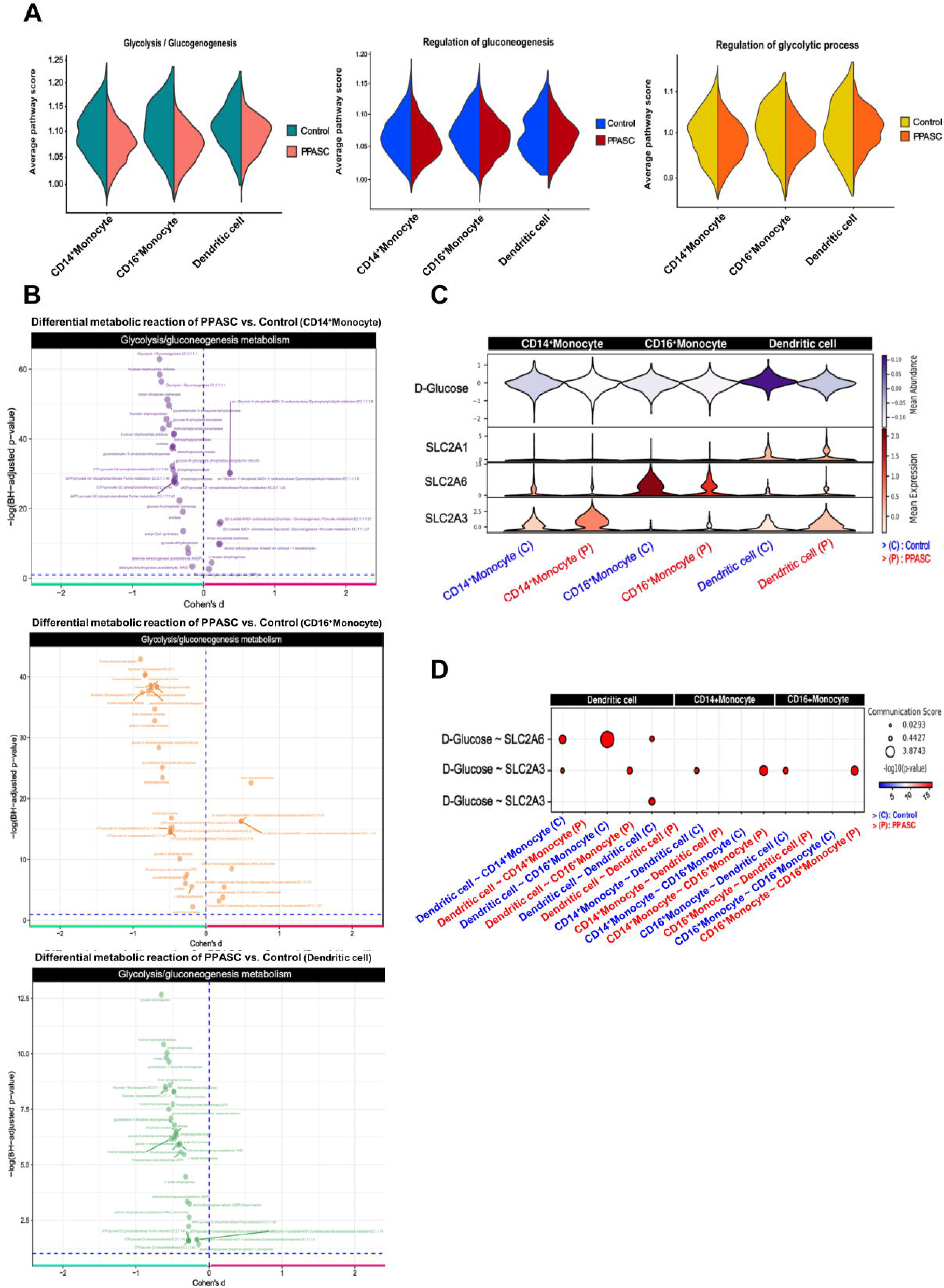
**A.** Gene signature of glycolysis-related pathways scored on average gene expression values in single cells. **B.** Metabolic reaction score in cells. **C.** Metabolite-sensor interactions between myeloid cells inferred from the total interactions of D-glucose and ATP. **D.** Inferred expression values of metabolites and sensors corresponding to D-Glucose and ATP.

## Discussion

Long-term perturbation of immune dysregulation and altered cytokine networks have been identified in COVID-19 convalescents and individuals with PASC tended to show more dysregulation of immune system (26, 48). This evidence support that altered immune system after recovery against SARS-CoV-2 infection contribute to development of clinical sequelae. While the epidemiological and clinical studies of PPASC individuals are relatively advanced, mechanistic insights into the pathophysiological underpinnings of this condition are still limited. In this study, we performed scRNA-seq of the PBMCs from PPASC patients and naïve control and an integrated analysis with published scRNA-seq of control dataset (65). We revealed that the enrichment of myeloid-specific cellular subsets, CD14^+^/CD16^+^ monocytes and dendritic cells showing increased pulmonary symptom/fibrosis signatures in PPASC. We further explored the cellular metabolism with analyzing metabolic reaction, metabolic pathway scoring, and cell-cell metabolite interaction from scRNA-seq data. To the best of our knowledge, this is the first time to show that PPASC individuals display metabolic alteration in MLCs with a skewing towards decreases in glycolysis and glucose utilization.

We recently reported persisting elevation of circulating monocytes in COVID-19 convalescents and the cell activation was negatively correlated with lung function in individuals with PPASC (66). Consistent with this finding, our scRNA-seq data demonstrates continued dysregulation of peripheral immune system, particularly MLCs after SARS-CoV-2 infection. In Ryan et al conducting scRNA-seq on PBMCs at 12-, 16-, and 24-weeks post-infection of SARS-CoV-2, they found that the blood transcriptome of convalescents was significantly perturbed compared to naïve controls (48). Indeed, transcriptional dysregulation persisted in individual with PASC compared to those who were completely recovered, until at least 24 weeks. Immunophenotyping result of PBMC demonstrated that specific subsets of CXCR3^+^ monocytes were increased in COVID-19 convalescents, but total monocytes were comparable between groups. None of the participants who were referred to the long-COVID clinic underwent PFTs, nor were they classified as pulmonary long-COVID. Because wide range of PASC symptoms mirrors the existence of different pathological subgroups, suggesting that increased monocytic populations observed within our study may be in relation to the pulmonary-specific immunopathology. In addition, pulmonary sequelae after SARS-CoV-2 infection PPASC individuals may have a selective immunological response that differ from the immunological response generated for other sequelae. Thus, more studies are essential for better understanding of the pathophysiologic mechanisms underlying diverse spectrums of PASC symptoms.

An evolving body of evidence suggests the importance of pro-inflammatory and pro-fibrotic monocytes and macrophages in PASC (68). In concert with our observed upregulation of gene expression related to pulmonary symptoms/fibrosis, suggesting that these cell types engage into sustaining fibrotic process that may fuel pulmonary sequelae. Interestingly, we also found that CD16^+^ gene module interaction networks showed the interactions with antigen-presentation genes. Antigen presentation on different monocyte subsets is well documented, with CD16^+^ intermediate and non-classical monocytes being the most prominent antigen presentation capable monocytes (69). In line with their findings, CD16^+^ monocytes were found to increase their expression of MHC molecules, further promoted by stimulation of Th1 skewing cytokines. We did not observe upregulation of any Th1 or Th2 skewing cytokines, suggesting this process is contributing towards our observed upregulation of MHC-2 molecules.

Recent study of plasma metabolomic profiles revealed dysregulation of metabolites with the enrichment of pathways for fatty acid metabolism and TCA cycles and high level of fatty acid metabolites (70). Advances in the field of immunometabolism have demonstrated strong associations of specific metabolic pathways with cellular phenotype. In addition, the metabolic and metabolite level of investigating within a cell are considered the most accurate thermometer for cellular function, as no processing or degradation (RNA/protein) is yet to occur. Although common residual symptoms of their study cohort were fatigue and brain fog, we observed similar trend of cellular metabolic alteration displaying the downregulation of gluconeogenesis/glycolysis pathways in MLCs. Furthermore, in the CD16^+^ M4 module genes suggesting a non-glycolysis-related metabolic signature, like ACSL4, a long-chain fatty acid ligase utilized for fatty acid beta-oxidation as well as OSBPL8, an oxysterol binding protein for intracellular lipid transport, were identified. These results suggest that the downregulation of gluconeogenesis/glycolysis pathways appeared to be associated with a compensatory upregulation of genes associated with fatty acid oxidation and amino acid-based metabolism. Future in-depth experimental work is required to understand whether metabolic reprogramming of monocytes after SARS-CoV-2 infection favors them to use fatty acid metabolism. Interestingly, mean abundance of ATP was relatively comparable between MLCs in both PPASC and controls, suggesting alternative metabolic pathways are pivotally compensatory in MLCs from PPASC, given the similar ATP production across the cell types.

A comprehensive review of Long-COVID disease, particularly chronic fatigue syndrome corroborates many of the findings observed from metabolic analyses of scRNA-seq of our study (71). Rewiring of cellular metabolism is directly linked to cellular function. Within monocyte populations, upregulation of glycolysis and glucose usage correlates with activation and differentiation towards an M1 i.e. pro-inflammatory monocyte/macrophage phenotype and functionality (72). This phenotype is defined by its pro-inflammatory cytokine production (IFN- y) and stimulation, while M2 relates to an anti-inflammatory and regulatory role (73). Use of fatty acid pathways and decreased glycolytic reliance suggests pro-resolution (M2)-like phenotype and functionality within these monocytes. Interestingly, this similar phenotype and reliance on fatty acid pathways has been observed in a model of SARS-CoV-2 vacciantion-induced myocarditis, specifally within monocytes (74). The authors reported that dysregulation of fatty acid and glycolysis pathways was associated with post-vaccination myocarditis, specifically within monocyte subsets isolated from PBMC (74). This same metabolic pattern is canonical during dengue virus infection of myeloid cells (75), but converse to the prototypical pattern of HIV infection of lymphocytes (76). In the future, it will be necessary to elucidate the role of monocytes immunometabolism in the context of PPASC and whether metabolic dysfunction is a predisposing risk factor for PPASC.

Our study was limited by a small sample size. One of our control participants was from a matched, publicly available single cell sequencing dataset. The comparator scRNA-seq data were further validated with a public dataset of other SARS-CoV-2 naïve single cell sequencing data to ensure continuity. Incorporation of further comparator groups from acute and recovered disease states could provide longitudinal insight into disease course and prognosis. Additionally, the use of other post-viral infection outcomes involving cellular perturbations or long-term metabolic shifts would aid in delineating SARS-CoV-2 similarities and differences to other conditions. Further studies in large-scale cohort studies, specifically targeting the pulmonary sequalae of long-COVID are needed in instrumental in developing our understanding of PPASC immunopathology.

## Conclusions

In conclusion, our analysis of scRNA-seq data reflecting pulmonary sequelae has revealed alteration of peripheral blood cells and associated gene signatures. Collectively, our data demonstrate an altered MLCs with specific skewing towards a pro-fibrotic and long-lived metabolic profiles in monocyte populations in PPASC. Such result provides important insights in mechanisms for dissecting the PASC symptoms and immunometabolism in the context of PPASC.

## Declarations

### Ethics approval and consent to participate

This study was approved by the Queen’s Medical Center Research & Institutional Review Committee RA-2020-053 and by the University of Hawaii Institutional Review Board 2020- 00406. Written informed consent was obtained from all participants and all assays were performed according to institutional guidelines and regulations.

### Consent for publication

Not applicable

### Availability of data and materials

The scRNA-seq sequences were deposited in NCBI GEO databases under the accession number GSE235938.

### Authors’ contributions

J.P. supervised the experimental approaches. B.J. collected and processed human blood samples.

G.D. analyzed PFT and clinical data collection. L.S.D. processed PBMC for scRNA-seq. and assisted in generating table. H.Y. and V.K. analyzed scRNA-seq data and H.Y. generated figures. D.C.C., V.R.K., and C.M.S. contributed to subject recruitment and sample collection. J.P., L.S.D, and H.Y wrote the draft of the manuscript. D.C.C, C.M.S., V.R.K., G.D., and Y.K., revised the manuscript. All the authors assisted in editing, provided critical review, and approved the final version of the submission.

---

**Supplementary Figure 1: A.** Total UMI counts were counted for each barcode to detect and remove empty droplets and droplets containing low amounts of RNA. **B.** Low-quality cells containing low UMI / feature counts and high mitochondrial gene expression were visualized by PCA to remove low-quality cells corresponding to outliers.

**Supplementary Table 1:** Demographics of naïve and PPASC participants. “–” denotes unavailable information.

## Acknowledgements

Authors thank the staff of the Hawaii Center for AIDS for participant recruitment, blood collection, and sample processing for the study during a particular challenging period. Dr. Alexandra Gurary at the Flow Cytometry Core, John A. Burns School of Medicine, University of Hawaii at Manoa who provided technical guidance for flow cytometry experiments. Dr. Karolina Peplowska at the Genomics and Bioinformatic Shared Resource, John A. Burns School of Medicine, The University of Hawaii Cancer Center who performed scRNA-seq. Authors are also grateful to the study participants.

## References

1. Wu Z, McGoogan JM. Characteristics of and Important Lessons from the Coronavirus Disease 2019 (Covid-19) Outbreak in China: Summary of a Report of 72 314 Cases from the Chinese Center for Disease Control and Prevention. JAMA (2020) 323(13):1239–42. doi: 10.1001/jama.2020.2648.

2. Raina SK, Kumar R. Identifying Role of Public Health and Primary Care as Disparate Entities in Current Health System. J Family Med Prim Care (2021) 10(10):3531–4. Epub 20211105. doi: 10.4103/jfmpc.jfmpc_1465_21.

3. Swerdlow DL, Finelli L. Preparation for Possible Sustained Transmission of 2019 Novel Coronavirus: Lessons from Previous Epidemics. JAMA (2020) 323(12):1129–30. doi: 10.1001/jama.2020.1960.

4. Pilkington V, Keestra SM, Hill A. Global Covid-19 Vaccine Inequity: Failures in the First Year of Distribution and Potential Solutions for the Future. Front Public Health (2022) 10:821117. Epub 20220307. doi: 10.3389/fpubh.2022.821117.

5. Tenforde MW, Self WH, Adams K, Gaglani M, Ginde AA, McNeal T, et al. Association between Mrna Vaccination and Covid-19 Hospitalization and Disease Severity. JAMA (2021) 326(20):2043–54. doi: 10.1001/jama.2021.19499.

6. Mohammed I, Nauman A, Paul P, Ganesan S, Chen KH, Jalil SMS, et al. The Efficacy and Effectiveness of the Covid-19 Vaccines in Reducing Infection, Severity, Hospitalization, and Mortality: A Systematic Review. Hum Vaccin Immunother (2022) 18(1):2027160. Epub 20220203. doi: 10.1080/21645515.2022.2027160.

7. Perlis RH, Santillana M, Ognyanova K, Safarpour A, Lunz Trujillo K, Simonson MD, et al. Prevalence and Correlates of Long Covid Symptoms among Us Adults. JAMA Netw Open (2022) 5(10):e2238804. Epub 20221003. doi: 10.1001/jamanetworkopen.2022.38804.

8. Zhou F, Tao M, Shang L, Liu Y, Pan G, Jin Y, et al. Assessment of Sequelae of Covid-19 Nearly 1 Year after Diagnosis. Front Med (Lausanne) (2021) 8:717194. Epub 20211123. doi: 10.3389/fmed.2021.717194.

9. Groff D, Sun A, Ssentongo AE, Ba DM, Parsons N, Poudel GR, et al. Short-Term and Long-Term Rates of Postacute Sequelae of Sars-Cov-2 Infection: A Systematic Review. JAMA Netw Open (2021) 4(10):e2128568. Epub 20211001. doi: 10.1001/jamanetworkopen.2021.28568.

10. Nalbandian A, Sehgal K, Gupta A, Madhavan MV, McGroder C, Stevens JS, et al. Post-Acute Covid-19 Syndrome. Nat Med (2021) 27(4):601–15. Epub 20210322. doi: 10.1038/s41591-021-01283-z.

11. Xie Y, Al-Aly Z. Risks and Burdens of Incident Diabetes in Long Covid: A Cohort Study. Lancet Diabetes Endocrinol (2022) 10(5):311–21. Epub 20220321. doi: 10.1016/S2213-8587(22)00044-4.

12. Xie Y, Xu E, Bowe B, Al-Aly Z. Long-Term Cardiovascular Outcomes of Covid-19. Nat Med (2022) 28(3):583–90. Epub 20220207. doi: 10.1038/s41591-022-01689-3.

13. Bowe B, Xie Y, Xu E, Al-Aly Z. Kidney Outcomes in Long Covid. J Am Soc Nephrol (2021) 32(11):2851–62. Epub 20210901. doi: 10.1681/ASN.2021060734.

14. O’Mahoney LL, Routen A, Gillies C, Ekezie W, Welford A, Zhang A, et al. The Prevalence and Long-Term Health Effects of Long Covid among Hospitalised and Non-Hospitalised Populations: A Systematic Review and Meta-Analysis. EClinicalMedicine (2023) 55:101762. Epub 20221201. doi: 10.1016/j.eclinm.2022.101762.

15. Global Burden of Disease Long CC, Wulf Hanson S, Abbafati C, Aerts JG, Al-Aly Z, Ashbaugh C, et al. Estimated Global Proportions of Individuals with Persistent Fatigue, Cognitive, and Respiratory Symptom Clusters Following Symptomatic Covid-19 in 2020 and 2021. JAMA (2022) 328(16):1604–15. doi: 10.1001/jama.2022.18931.

16. Chen C, Haupert SR, Zimmermann L, Shi X, Fritsche LG, Mukherjee B. Global Prevalence of Post-Coronavirus Disease 2019 (Covid-19) Condition or Long Covid: A Meta-Analysis and Systematic Review. J Infect Dis (2022) 226(9):1593–607. doi: 10.1093/infdis/jiac136.

17. McFann K, Baxter BA, LaVergne SM, Stromberg S, Berry K, Tipton M, et al. Quality of Life (Qol) Is Reduced in Those with Severe Covid-19 Disease, Post-Acute Sequelae of Covid-19, and Hospitalization in United States Adults from Northern Colorado. Int J Environ Res Public Health (2021) 18(21). Epub 20211021. doi: 10.3390/ijerph182111048.

18. Shah R, Ali FM, Nixon SJ, Ingram JR, Salek SM, Finlay AY. Measuring the Impact of Covid-19 on the Quality of Life of the Survivors, Partners and Family Members: A Cross-Sectional International Online Survey. BMJ Open (2021) 11(5):e047680. Epub 20210525. doi: 10.1136/bmjopen-2020-047680.

19. Tak CR. The Health Impact of Long Covid: A Cross-Sectional Examination of Health-Related Quality of Life, Disability, and Health Status among Individuals with Self-Reported Post-Acute Sequelae of Sars Cov-2 Infection at Various Points of Recovery. J Patient Rep Outcomes (2023) 7(1):31. Epub 20230321. doi: 10.1186/s41687-023-00572-0.

20. Tsampasian V, Elghazaly H, Chattopadhyay R, Debski M, Naing TKP, Garg P, et al. Risk Factors Associated with Post-Covid-19 Condition: A Systematic Review and Meta-Analysis. JAMA Intern Med (2023). Epub 20230323. doi: 10.1001/jamainternmed.2023.0750.

21. Lukkahatai N, Rodney T, Ling C, Daniel B, Han HR. Long Covid in the Context of Social Determinants of Health. Front Public Health (2023) 11:1098443. Epub 20230328. doi: 10.3389/fpubh.2023.1098443.

22. Khullar D, Zhang Y, Zang C, Xu Z, Wang F, Weiner MG, et al. Racial/Ethnic Disparities in Post-Acute Sequelae of Sars-Cov-2 Infection in New York: An Ehr-Based Cohort Study from the Recover Program. J Gen Intern Med (2023) 38(5):1127–36. Epub 20230216. doi: 10.1007/s11606-022-07997-1.

23. Xie Y, Bowe B, Al-Aly Z. Burdens of Post-Acute Sequelae of Covid-19 by Severity of Acute Infection, Demographics and Health Status. Nat Commun (2021) 12(1):6571. Epub 20211112. doi: 10.1038/s41467-021-26513-3.

24. Bai F, Tomasoni D, Falcinella C, Barbanotti D, Castoldi R, Mule G, et al. Female Gender Is Associated with Long Covid Syndrome: A Prospective Cohort Study. Clin Microbiol Infect (2022) 28(4):611 e9- e16. Epub 20211109. doi: 10.1016/j.cmi.2021.11.002.

25. Klein J, Wood J, Jaycox J, Lu P, Dhodapkar RM, Gehlhausen JR, et al. Distinguishing Features of Long Covid Identified through Immune Profiling. medRxiv (2022). Epub 20220810. doi: 10.1101/2022.08.09.22278592.

26. Phetsouphanh C, Darley DR, Wilson DB, Howe A, Munier CML, Patel SK, et al. Immunological Dysfunction Persists for 8 Months Following Initial Mild-to-Moderate Sars-Cov- 2 Infection. Nat Immunol (2022) 23(2):210–6. Epub 20220113. doi: 10.1038/s41590-021-01113-x.

27. Cui L, Fang Z, De Souza CM, Lerbs T, Guan Y, Li I, et al. Innate Immune Cell Activation Causes Lung Fibrosis in a Humanized Model of Long Covid. Proc Natl Acad Sci U S A (2023) 120(10):e2217199120. Epub 20230227. doi: 10.1073/pnas.2217199120.

28. Cheon IS, Li C, Son YM, Goplen NP, Wu Y, Cassmann T, et al. Immune Signatures Underlying Post-Acute Covid-19 Lung Sequelae. Sci Immunol (2021) 6(65):eabk1741. Epub 20211112. doi: 10.1126/sciimmunol.abk1741.

29. Bodansky A, Wang CY, Saxena A, Mitchell A, Takahashi S, Anglin K, et al. Autoantigen Profiling Reveals a Shared Post-Covid Signature in Fully Recovered and Long Covid Patients. medRxiv (2023). Epub 20230209. doi: 10.1101/2023.02.06.23285532.

30. Peluso MJ, Lu S, Tang AF, Durstenfeld MS, Ho HE, Goldberg SA, et al. Markers of Immune Activation and Inflammation in Individuals with Postacute Sequelae of Severe Acute Respiratory Syndrome Coronavirus 2 Infection. J Infect Dis (2021) 224(11):1839–48. doi: 10.1093/infdis/jiab490.

31. Schultheiss C, Willscher E, Paschold L, Gottschick C, Klee B, Henkes SS, et al. The Il- 1beta, Il-6, and Tnf Cytokine Triad Is Associated with Post-Acute Sequelae of Covid-19. Cell Rep Med (2022) 3(6):100663. doi: 10.1016/j.xcrm.2022.100663.

32. Espin E, Yang C, Shannon CP, Assadian S, He D, Tebbutt SJ. Cellular and Molecular Biomarkers of Long Covid: A Scoping Review. EBioMedicine (2023) 91:104552. Epub 20230408. doi: 10.1016/j.ebiom.2023.104552.

33. Thomas T, Stefanoni D, Reisz JA, Nemkov T, Bertolone L, Francis RO, et al. Covid-19 Infection Alters Kynurenine and Fatty Acid Metabolism, Correlating with Il-6 Levels and Renal Status. *JCI Insight* (2020) 5(14). Epub 20200723. doi: 10.1172/jci.insight.140327.

34. Feingold KR. The Bidirectional Link between Hdl and Covid-19 Infections. J Lipid Res (2021) 62:100067. Epub 20210317. doi: 10.1016/j.jlr.2021.100067.

35. Scherer PE, Kirwan JP, Rosen CJ. Post-Acute Sequelae of Covid-19: A Metabolic Perspective. Elife (2022) 11. Epub 20220323. doi: 10.7554/eLife.78200.

36. Dias SSG, Soares VC, Ferreira AC, Sacramento CQ, Fintelman-Rodrigues N, Temerozo JR, et al. Lipid Droplets Fuel Sars-Cov-2 Replication and Production of Inflammatory Mediators. PLoS Pathog (2020) 16(12):e1009127. Epub 20201216. doi: 10.1371/journal.ppat.1009127.

37. Nardacci R, Colavita F, Castilletti C, Lapa D, Matusali G, Meschi S, et al. Evidences for Lipid Involvement in Sars-Cov-2 Cytopathogenesis. Cell Death Dis (2021) 12(3):263. Epub 20210312. doi: 10.1038/s41419-021-03527-9.

38. Grootemaat AE, van der Niet S, Scholl ER, Roos E, Schurink B, Bugiani M, et al. Lipid and Nucleocapsid N-Protein Accumulation in Covid-19 Patient Lung and Infected Cells. Microbiol Spectr (2022) 10(1):e0127121. Epub 20220216. doi: 10.1128/spectrum.01271-21.

39. Codo AC, Davanzo GG, Monteiro LB, de Souza GF, Muraro SP, Virgilio-da-Silva JV, et al. Elevated Glucose Levels Favor Sars-Cov-2 Infection and Monocyte Response through a Hif- 1alpha/Glycolysis-Dependent Axis. Cell Metab (2020) 32(3):437–46 e5. Epub 20200717. doi: 10.1016/j.cmet.2020.07.007.

40. Gurshaney S, Morales-Alvarez A, Ezhakunnel K, Manalo A, Huynh TH, Abe JI, et al. Metabolic Dysregulation Impairs Lymphocyte Function During Severe Sars-Cov-2 Infection. Commun Biol (2023) 6(1):374. Epub 20230407. doi: 10.1038/s42003-023-04730-4.

41. Kovarik JJ, Bileck A, Hagn G, Meier-Menches SM, Frey T, Kaempf A, et al. A Multi- Omics Based Anti-Inflammatory Immune Signature Characterizes Long Covid-19 Syndrome. iScience (2023) 26(1):105717. Epub 20221205. doi: 10.1016/j.isci.2022.105717.

42. Gaebler C, Wang Z, Lorenzi JCC, Muecksch F, Finkin S, Tokuyama M, et al. Evolution of Antibody Immunity to Sars-Cov-2. Nature (2021) 591(7851):639–44. Epub 20210118. doi: 10.1038/s41586-021-03207-w.

43. Stein SR, Ramelli SC, Grazioli A, Chung JY, Singh M, Yinda CK, et al. Sars-Cov-2 Infection and Persistence in the Human Body and Brain at Autopsy. Nature (2022) 612(7941):758-63. Epub 20221214. doi: 10.1038/s41586-022-05542-y.

44. Patterson BK, Francisco EB, Yogendra R, Long E, Pise A, Rodrigues H, et al. Persistence of Sars Cov-2 S1 Protein in Cd16+ Monocytes in Post-Acute Sequelae of Covid-19 (Pasc) up to 15 Months Post-Infection. Front Immunol (2021) 12:746021. Epub 20220110. doi: 10.3389/fimmu.2021.746021.

45. Cohn LB, Chomont N, Deeks SG. The Biology of the Hiv-1 Latent Reservoir and Implications for Cure Strategies. Cell Host Microbe (2020) 27(4):519–30. doi: 10.1016/j.chom.2020.03.014.

46. Hendricks CM, Cordeiro T, Gomes AP, Stevenson M. The Interplay of Hiv-1 and Macrophages in Viral Persistence. Front Microbiol (2021) 12:646447. Epub 20210407. doi: 10.3389/fmicb.2021.646447.

47. Wu VH, Nordin JML, Nguyen S, Joy J, Mampe F, Del Rio Estrada PM, et al. Profound Phenotypic and Epigenetic Heterogeneity of the Hiv-1-Infected Cd4(+) T Cell Reservoir. Nat Immunol (2023) 24(2):359–70. Epub 20221219. doi: 10.1038/s41590-022-01371-3.

48. Ryan FJ, Hope CM, Masavuli MG, Lynn MA, Mekonnen ZA, Yeow AEL, et al. Long-Term Perturbation of the Peripheral Immune System Months after Sars-Cov-2 Infection. BMC Med (2022) 20(1):26. Epub 20220114. doi: 10.1186/s12916-021-02228-6.

49. Su Y, Yuan D, Chen DG, Ng RH, Wang K, Choi J, et al. Multiple Early Factors Anticipate Post-Acute Covid-19 Sequelae. Cell (2022) 185(5):881–95 e20. Epub 20220125. doi: 10.1016/j.cell.2022.01.014.

50. Graham BL, Brusasco V, Burgos F, Cooper BG, Jensen R, Kendrick A, et al. 2017 Ers/Ats Standards for Single-Breath Carbon Monoxide Uptake in the Lung. Eur Respir J (2017) 49(1). Epub 20170103. doi: 10.1183/13993003.00016-2016.

51. Zheng GX, Terry JM, Belgrader P, Ryvkin P, Bent ZW, Wilson R, et al. Massively Parallel Digital Transcriptional Profiling of Single Cells. Nat Commun (2017) 8:14049. Epub 20170116. doi: 10.1038/ncomms14049.

52. Lun ATL, Riesenfeld S, Andrews T, Dao TP, Gomes T, participants in the 1st Human Cell Atlas J, et al. Emptydrops: Distinguishing Cells from Empty Droplets in Droplet-Based Single-Cell Rna Sequencing Data. Genome Biol (2019) 20(1):63. Epub 20190322. doi: 10.1186/s13059-019-1662-y.

53. McCarthy DJ, Campbell KR, Lun AT, Wills QF. Scater: Pre-Processing, Quality Control, Normalization and Visualization of Single-Cell Rna-Seq Data in R. Bioinformatics (2017) 33(8):1179–86. doi: 10.1093/bioinformatics/btw777.

54. Lun AT, McCarthy DJ, Marioni JC. A Step-by-Step Workflow for Low-Level Analysis of Single-Cell Rna-Seq Data with Bioconductor. F1000Res (2016) 5:2122. Epub 20160831. doi: 10.12688/f1000research.9501.2.

55. Korsunsky I, Millard N, Fan J, Slowikowski K, Zhang F, Wei K, et al. Fast, Sensitive and Accurate Integration of Single-Cell Data with Harmony. Nat Methods (2019) 16(12):1289–96. Epub 20191118. doi: 10.1038/s41592-019-0619-0.

56. Hu C, Li T, Xu Y, Zhang X, Li F, Bai J, et al. Cellmarker 2.0: An Updated Database of Manually Curated Cell Markers in Human/Mouse and Web Tools Based on Scrna-Seq Data. Nucleic Acids Res (2023) 51(D1):D870–D6. doi: 10.1093/nar/gkac947.

57. Miller SA, Policastro RA, Sriramkumar S, Lai T, Huntington TD, Ladaika CA, et al. Lsd1 and Aberrant DNA Methylation Mediate Persistence of Enteroendocrine Progenitors That Support Braf-Mutant Colorectal Cancer. Cancer Res (2021) 81(14):3791–805. Epub 20210525. doi: 10.1158/0008-5472.CAN-20-3562.

58. Finak G, McDavid A, Yajima M, Deng J, Gersuk V, Shalek AK, et al. Mast: A Flexible Statistical Framework for Assessing Transcriptional Changes and Characterizing Heterogeneity in Single-Cell Rna Sequencing Data. Genome Biol (2015) 16:278. Epub 20151210. doi: 10.1186/s13059-015-0844-5.

59. Liberzon A, Subramanian A, Pinchback R, Thorvaldsdottir H, Tamayo P, Mesirov JP. Molecular Signatures Database (Msigdb) 3.0. Bioinformatics (2011) 27(12):1739–40. Epub 20110505. doi: 10.1093/bioinformatics/btr260.

60. Zheng R, Zhang Y, Tsuji T, Zhang L, Tseng Y-H, Chen K. Mebocost: Metabolic Cell-Cell Communication Modeling by Single Cell Transcriptome. bioRxiv (2022):2022.05.30.494067. doi: 10.1101/2022.05.30.494067.

61. Jin S, Guerrero-Juarez CF, Zhang L, Chang I, Ramos R, Kuan CH, et al. Inference and Analysis of Cell-Cell Communication Using Cellchat. Nat Commun (2021) 12(1):1088. Epub 20210217. doi: 10.1038/s41467-021-21246-9.

62. Robinson MD, McCarthy DJ, Smyth GK. Edger: A Bioconductor Package for Differential Expression Analysis of Digital Gene Expression Data. Bioinformatics (2010) 26(1):139–40. Epub 20091111. doi: 10.1093/bioinformatics/btp616.

63. Morabito S, Reese F, Rahimzadeh N, Miyoshi E, Swarup V. High Dimensional Co-Expression Networks Enable Discovery of Transcriptomic Drivers in Complex Biological Systems. bioRxiv (2022):2022.09.22.509094. doi: 10.1101/2022.09.22.509094.

64. Chen EY, Tan CM, Kou Y, Duan Q, Wang Z, Meirelles GV, et al. Enrichr: Interactive and Collaborative Html5 Gene List Enrichment Analysis Tool. BMC Bioinformatics (2013) 14:128. Epub 20130415. doi: 10.1186/1471-2105-14-128.

65. Lee JS, Park S, Jeong HW, Ahn JY, Choi SJ, Lee H, et al. Immunophenotyping of Covid- 19 and Influenza Highlights the Role of Type I Interferons in Development of Severe Covid-19. Sci Immunol (2020) 5(49). doi: 10.1126/sciimmunol.abd1554.

66. Park J, Dean LS, Jiyarom B, Gangcuangco LM, Shah P, Awamura T, et al. Elevated Circulating Monocytes and Monocyte Activation in Covid-19 Convalescent Individuals. Front Immunol (2023) 14:1151780. Epub 20230403. doi: 10.3389/fimmu.2023.1151780.

67. Gilmcher MJ, Katz EP. The Organization of Collagen in Bone: The Role of Noncovalent Bonds in the Relative Insolubility of Bone Collagen. J Ultrastruct Res (1965) 12(5):705–29. doi: 10.1016/s0022-5320(65)80057-0.

68. Kamp JC, Werlein C, Plucinski EKJ, Neubert L, Welte T, Lee PD, et al. Novel Insight into Pulmonary Fibrosis and Long Covid. Am J Respir Crit Care Med (2023) 207(8):1105–7. doi: 10.1164/rccm.202212-2314LE.

69. Lee J, Tam H, Adler L, Ilstad-Minnihan A, Macaubas C, Mellins ED. The Mhc Class Ii Antigen Presentation Pathway in Human Monocytes Differs by Subset and Is Regulated by Cytokines. PLoS One (2017) 12(8):e0183594. Epub 20170823. doi: 10.1371/journal.pone.0183594.

70. Guntur VP, Nemkov T, de Boer E, Mohning MP, Baraghoshi D, Cendali FI, et al. Signatures of Mitochondrial Dysfunction and Impaired Fatty Acid Metabolism in Plasma of Patients with Post-Acute Sequelae of Covid-19 (Pasc). Metabolites (2022) 12(11). Epub 20221026. doi: 10.3390/metabo12111026.

71. Davis HE, McCorkell L, Vogel JM, Topol EJ. Long Covid: Major Findings, Mechanisms and Recommendations. Nat Rev Microbiol (2023) 21(3):133–46. Epub 20230113. doi: 10.1038/s41579-022-00846-2.

72. Zhu X, Meyers A, Long D, Ingram B, Liu T, Yoza BK, et al. Frontline Science: Monocytes Sequentially Rewire Metabolism and Bioenergetics During an Acute Inflammatory Response. J Leukoc Biol (2019) 105(2):215–28. Epub 20190111. doi: 10.1002/JLB.3HI0918-373R.

73. Stunault MI, Bories G, Guinamard RR, Ivanov S. Metabolism Plays a Key Role During Macrophage Activation. Mediators Inflamm (2018) 2018:2426138. Epub 20181210. doi: 10.1155/2018/2426138.

74. Hwang N, Huh Y, Bu S, Seo KJ, Kwon SH, Kim JW, et al. Single-Cell Sequencing of Pbmc Characterizes the Altered Transcriptomic Landscape of Classical Monocytes in Bnt162b2- Induced Myocarditis. Front Immunol (2022) 13:979188. Epub 20220926. doi: 10.3389/fimmu.2022.979188.

75. Thyrsted J, Holm CK. Virus-Induced Metabolic Reprogramming and Innate Sensing Hereof by the Infected Host. Curr Opin Biotechnol (2021) 68:44–50. Epub 20201025. doi: 10.1016/j.copbio.2020.10.004.

76. Butterfield TR, Landay AL, Anzinger JJ. Dysfunctional Immunometabolism in Hiv Infection: Contributing Factors and Implications for Age-Related Comorbid Diseases. Curr HIV/AIDS Rep (2020) 17(2):125–37. doi: 10.1007/s11904-020-00484-4.

