## Supplementary Table 1 for "Single-cell RNA sequencing reveals characteristics of myeloid cells in pulmonary post-acute sequelae of SARS-CoV-2"

|  | **Control 1** | **PPASC 1** | **PPASC 2** |
| --- | --- | --- | --- |
| **Age** | 61 | 52 | 60 |
| **Ethnicity** | Chinese | Japanese | Japanese |
| **BMI** | - | 29.3 | 26.2 |
| **History of Asthma** | No | Yes | No |
| **History of COPD** | No | No | Yes |
| **Ever Smoked** | No | No | No |
| **Hospitalized for COVID-19** | No | Yes | Yes |
| **Date Enrolled Post-Acute Infection** | 0 | 9 | 5 |
| **Fully Vaccinated against SARS-CoV-2** | - | Yes | Yes |
| **WBC** | - | 5.86 | 10.38 |
| **Monocyte Count** | - | 9.9 | 7.5 |
| **Neutrophil Count** | - | 2.36 | 7.87 |
| **Platelet Count** | - | 264 | 221 |
| **Lymphocyte Count** | - | 2.61 | 1.4 |
| **D-Dimer** | - | 0.41 | 0.48 |
| **CRP** | - | 3.0 | <3.0 |
| **DLCO-c%** | - | 66.4 | 39.2 |

Demographics of naïve and PPASC participants. “–” denotes unavailable information.
