## Supplementary figures and images for "Single-cell RNA sequencing reveals characteristics of myeloid cells in pulmonary post-acute sequelae of SARS-CoV-2"

### Supplementary Figure 1

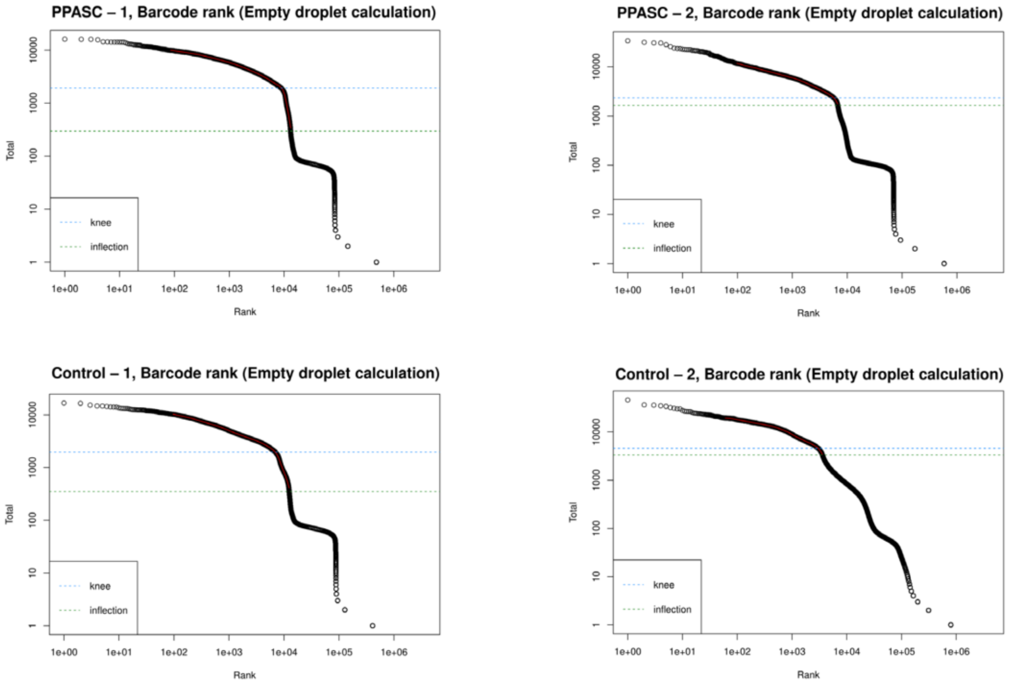

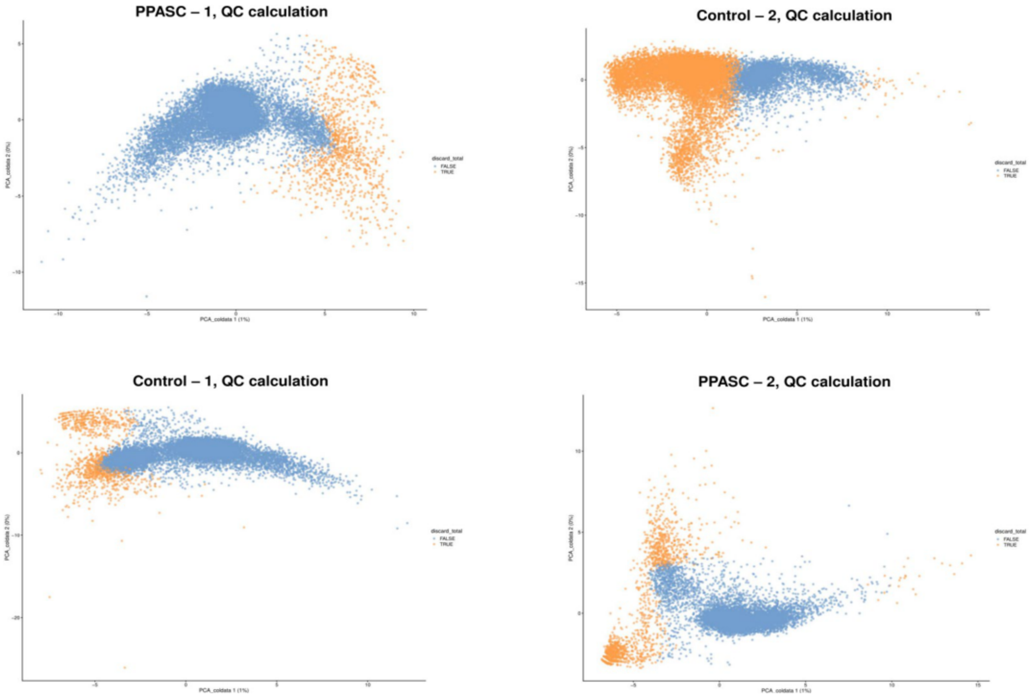
